## Supplementary material for "On the analysis of genetic association with long-read sequencing data": S1 Text

### S4 Text: Relationships between RoP, saturated test and 4 df interaction test

| Diplotypes | Additive | | Dominance | | Interaction and phase terms | | | | | Phasetypes | | | |
|---|---|---|---|---|---|---|---|---|---|---|---|---|---|
| | $G_A$ | $G_B$ | $D_A$ | $D_B$ | $G_A \cdot G_B$ | $G_A \cdot D_B$ | $D_A \cdot G_B$ | $D_A \cdot D_B$ | $V$ | $P_{cis}$ | $D_{cis}$ | $P_{trans}$ | $D_{trans}$ |
| ab/ab | 0 | 0 | 0 | 0 | 0 | 0 | 0 | 0 | 0 | 0 | 0 | 0 | 0 |
| Ab/ab | 1 | 0 | 1 | 0 | 0 | 0 | 0 | 0 | 0 | 0 | 0 | 1 | 1 |
| aB/ab | 0 | 1 | 0 | 1 | 0 | 0 | 0 | 0 | 0 | 0 | 1 | 0 | 0 |
| AB/ab | 1 | 1 | 1 | 1 | 1 | 1 | 1 | 1 | 1 | 1 | 1 | 0 | 1 |
| Ab/aB | 1 | 1 | 1 | 1 | 0 | 1 | 1 | 1 | 0 | 0 | 0 | 1 | 0 |
| Ab/Ab | 2 | 0 | 0 | 0 | 0 | 0 | 0 | 0 | 0 | 0 | 0 | 0 | 0 |
| aB/aB | 0 | 2 | 0 | 0 | 0 | 0 | 0 | 0 | 0 | 0 | 0 | 0 | 0 |
| AB/aB | 1 | 2 | 1 | 0 | 1 | 0 | 2 | 0 | 0 | 1 | 1 | 1 | 0 |
| AB/Ab | 2 | 1 | 0 | 1 | 1 | 2 | 0 | 0 | 0 | 1 | 0 | 1 | 1 |
| AB/AB | 2 | 2 | 0 | 0 | 4 | 0 | 0 | 0 | 0 | 2 | 0 | 2 | 0 |

Table 1: Genotypes, interaction terms, the phase term and the additive and dominance phasetypes of two bialleleic loci with alleles A,a and B,b. The phasetypes $P_{cis}$ and $P_{trans}$ are coded with respect to allele A and B.

The additive *cis* and *trans* relationships can be written as the linear combinations of the phase term and the four genotype interaction terms as in Table 1:

$$P_{cis} = V + \frac{1}{2} \cdot (-G_A \cdot D_B + G_A \cdot G_B - D_A \cdot G_B + D_A \cdot D_B)$$

$$P_{trans} = -V + \frac{1}{2} \cdot (G_A \cdot D_B + G_A \cdot G_B + D_A \cdot G_B - D_A \cdot D_B)$$

Therefore, the saturated model, which tests the epistasis and phase effects by the 5 df test $H_0 : \beta_{G_A G_B} = \beta_{G_A D_B} = \beta_{D_A G_B} = \beta_{D_A D_B} = \beta_V = 0$, can fully capture the additive phase effects. However, the additional degrees of freedom reduces its power compared to the 1 df RoP tests

The saturated test cannot effectively distinguish *cis* effects from *trans* effects. In particular, the direction of $\beta_V$ relies on both *cis* and *trans* contribution as well as the reference alleles for the phasetypes. Below we summarize the directions of $\beta_V$:

| Contributing phase relationships | Direction of $\beta_V$ |
|---|---|
| $Cis_{AB}$ | + |
| $Cis_{Ab}$ | - |
| $Cis_{aB}$ | - |
| $Cis_{ab}$ | + |
| $Trans_{AB}$ | - |
| $Trans_{Ab}$ | + |
| $Trans_{aB}$ | + |
| $Trans_{ab}$ | - |

The recessive inheritance patterns of *cis* and *trans* relationships are identical, i.e., $I_{cis=2} = I_{trans=2}$, where $I$ is the indicator variable that equals 1 when $Cis_{AB} = 2$ or $Trans_{AB} = 2$, respectively. Both *cis* and *trans* relationships can represent recessive effects with their additive and dominance phasetypes:

$$I_{Cis=2} = \frac{1}{2}(Cis_{AB} - D_{cis}) = \frac{1}{2}(Trans_{AB} - D_{trans})$$

therefore, unlike the single locus analysis with genotypes, the power of RoP to detect recessive *cis* effects cannot be improved by simply including the dominance terms for *cis* and *trans* relationships and testing $H_0 : \beta_{P_{cis}} = \beta_{D_{cis}} = 0$, as the recessive *cis* effects can also well explained by the remaining additive and dominance terms of *trans* relationships.

Instead, the recessive *cis* effect can be modelled by the 4 interaction terms

$$I_{Cis=2} = \frac{1}{4} \cdot (-G_A \cdot D_B + G_A \cdot G_B - D_A \cdot G_B + D_A \cdot D_B)$$

and the 4 df interaction test has the best performance in detecting recessive *cis* effects, even it doesn't include phase information.
