## Supplementary figures and images for "On the analysis of genetic association with long-read sequencing data"

### S1 Fig

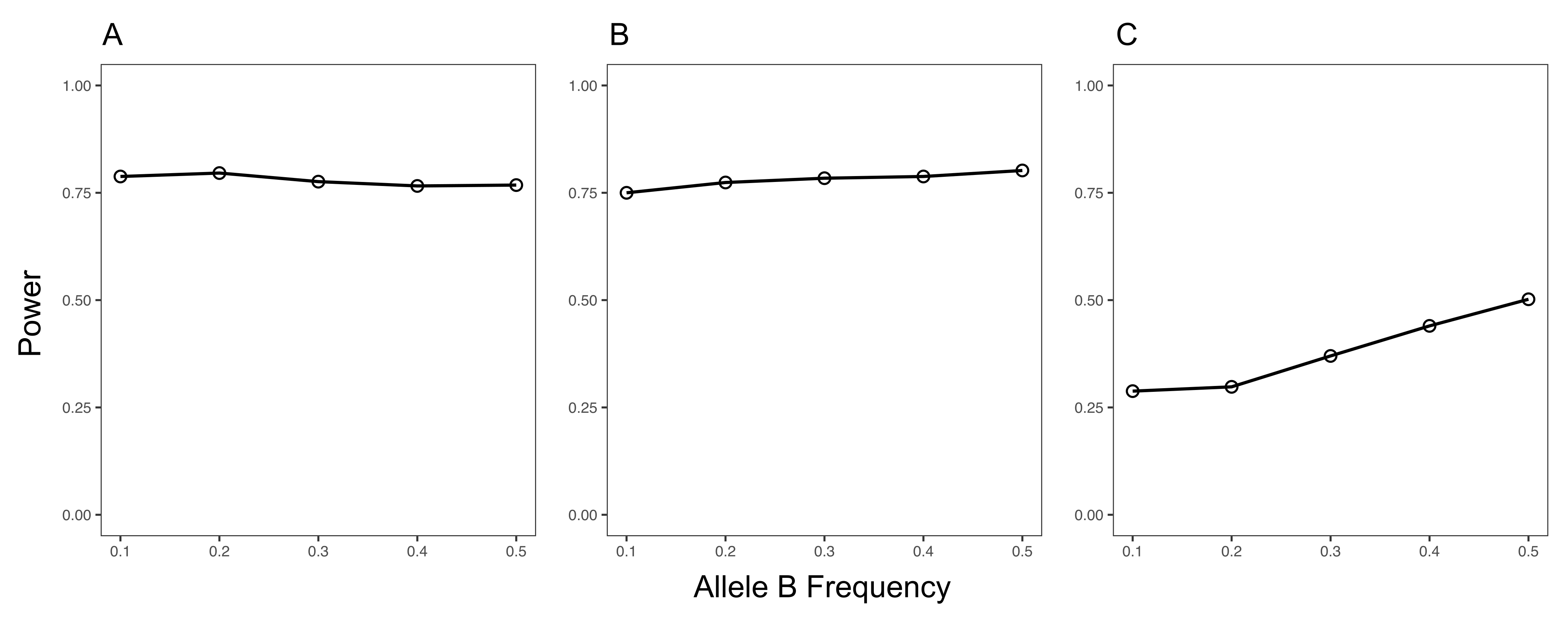

### S2 Fig

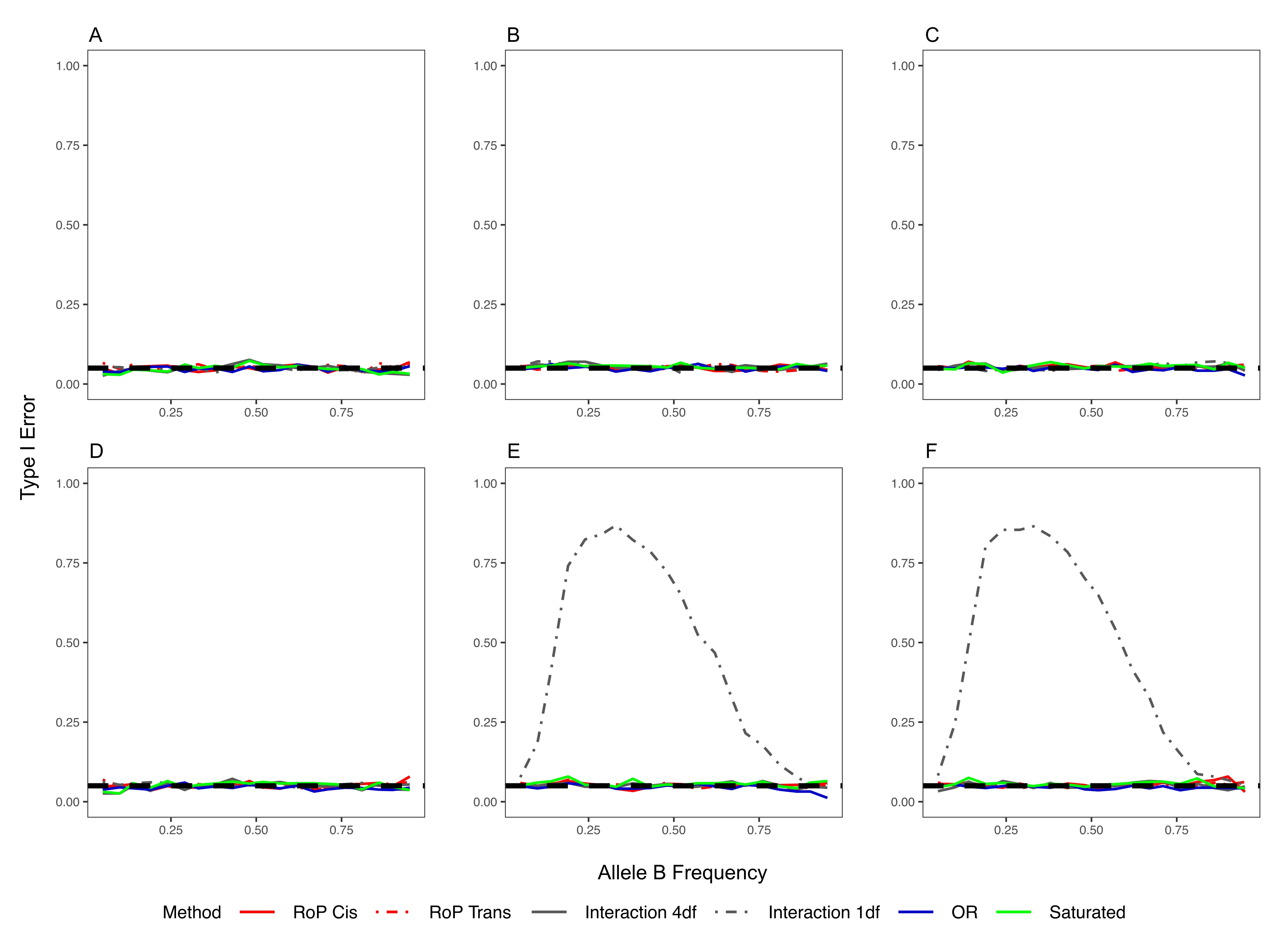

### S3 Fig

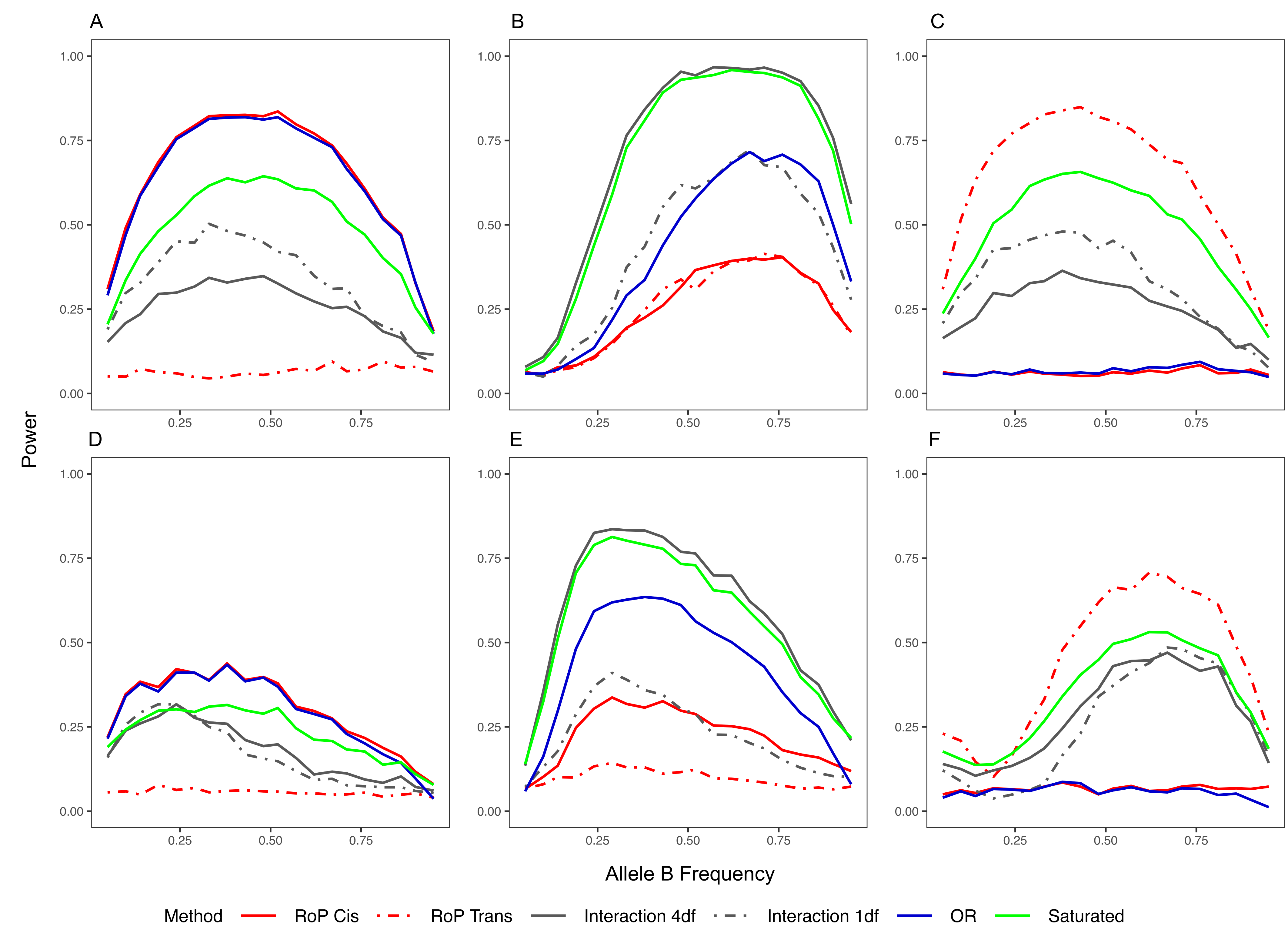

### S4 Fig

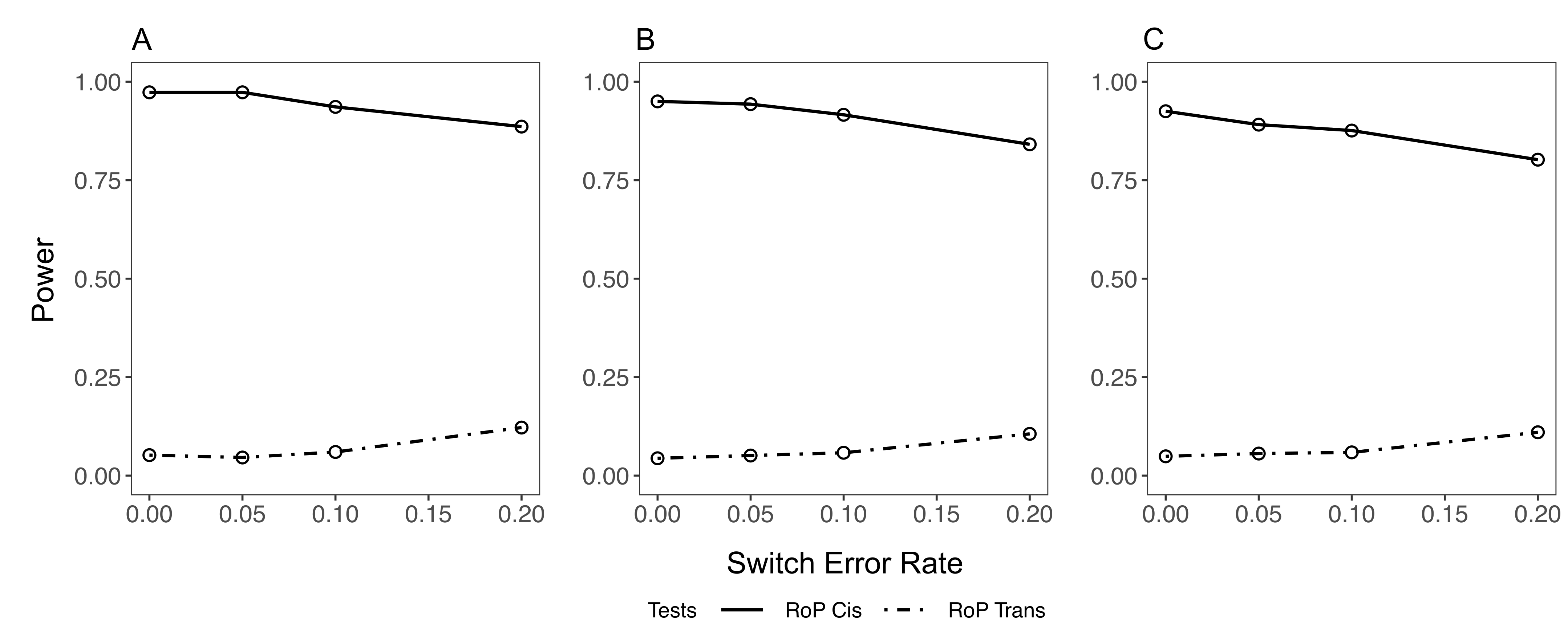
