## Supplementary material for "On the analysis of genetic association with long-read sequencing data": S3 Text

### S3 Text: The haploypte frequency odds-ratio (OR) approximates additive *cis* effects

The haplotype frequency OR test measures the interaction effects between two biallelic variants by the "odds-ratio" :

$$log(OR) = log\frac{h_{AB}h_{ab}}{h_{aB}h_{Ab}} - log\frac{(1-h_{AB})(1-h_{ab})}{(1-h_{aB})(1-h_{Ab})}$$

$h_{ij}$ is the haplotype penetrance, defined as:

$$h_{ij} = \phi_{AB}f_{ijAB} + \phi_{Ab}f_{ijAb} + \phi_{aB}f_{ijaB} + \phi_{ab}f_{ijab}$$

and $f_{jikl}$ is the penetrance for diplotype ij/kl, which is the probability of having the disease if one carries the diplotype ij/kl.

Ueki et al, 2012. showed that $log(OR)$ approximates the interaction effect $\beta_{G_A \cdot G_B}$ under the rare disease assumption. Here, we show that $log(OR)$ approximates $\beta_{cis}$ following the same approach.

For an additive *cis* effect, we consider the following logistic model for a case-control outcome:

$$logit(p) = \alpha + \beta \cdot Cis$$

the penetrance for each diplotype $f_{ijkl}$ can be written as

| $f_{ijkl}$ | AB | Ab | aB | ab |
|---|---|---|---|---|
| AB | $\frac{e^{\alpha+2\beta}}{1+e^{\alpha+2\beta}}$ | $\frac{e^{\alpha+\beta}}{1+e^{\alpha+\beta}}$ | $\frac{e^{\alpha+\beta}}{1+e^{\alpha+\beta}}$ | $\frac{e^{\alpha+\beta}}{1+e^{\alpha+\beta}}$ |
| Ab | $\frac{e^{\alpha+\beta}}{1+e^{\alpha+\beta}}$ | $\frac{e^{\alpha}}{1+e^{\alpha}}$ | $\frac{e^{\alpha}}{1+e^{\alpha}}$ | $\frac{e^{\alpha}}{1+e^{\alpha}}$ |
| aB | $\frac{e^{\alpha+\beta}}{1+e^{\alpha+\beta}}$ | $\frac{e^{\alpha}}{1+e^{\alpha}}$ | $\frac{e^{\alpha}}{1+e^{\alpha}}$ | $\frac{e^{\alpha}}{1+e^{\alpha}}$ |
| ab | $\frac{e^{\alpha+\beta}}{1+e^{\alpha+\beta}}$ | $\frac{e^{\alpha}}{1+e^{\alpha}}$ | $\frac{e^{\alpha}}{1+e^{\alpha}}$ | $\frac{e^{\alpha}}{1+e^{\alpha}}$ |

Under the rare disease assumption, the diplotype penetrance can be simplified to

| $f_{ijkl}$ | AB | Ab | aB | ab |
|---|---|---|---|---|
| AB | $e^{\alpha+2\beta}$ | $e^{\alpha+\beta}$ | $e^{\alpha+\beta}$ | $e^{\alpha+\beta}$ |
| Ab | $e^{\alpha+\beta}$ | $e^{\alpha}$ | $e^{\alpha}$ | $e^{\alpha}$ |
| aB | $e^{\alpha+\beta}$ | $e^{\alpha}$ | $e^{\alpha}$ | $e^{\alpha}$ |
| ab | $e^{\alpha+\beta}$ | $e^{\alpha}$ | $e^{\alpha}$ | $e^{\alpha}$ |

and we also have

$$log\frac{(1-h_{AB})(1-h_{ab})}{(1-h_{aB})(1-h_{Ab})} \approx 1$$

Then

$$\begin{aligned}
log(OR) &= log\frac{h_{AB}h_{ab}}{h_{aB}h_{Ab}} \\
&= log\frac{(\phi_{AB}f_{ABAB} + \phi_{Ab}f_{ABAb} + \phi_{aB}f_{ABaB} + \phi_{ab}f_{ABab})(\phi_{AB}f_{abAB} + \phi_{Ab}f_{abABAb} + \phi_{aB}f_{abABaB} + \phi_{ab}f_{abABab})}{(\phi_{AB}f_{AbAB} + \phi_{Ab}f_{AbAb} + \phi_{aB}f_{AbaB} + \phi_{ab}f_{Abab})(\phi_{AB}f_{aBAB} + \phi_{Ab}f_{aBAb} + \phi_{aB}f_{aBaB} + \phi_{ab}f_{aBab})} \\
&= log\frac{e^{\alpha}(\phi_{AB}e^{2\beta} + \phi_{Ab}e^{\beta} + \phi_{aB}e^{\beta} + \phi_{ab}e^{\beta})e^{\alpha}(\phi_{AB}e^{\beta} + \phi_{Ab} + \phi_{aB} + \phi_{ab})}{e^{\alpha}(\phi_{AB}e^{\beta} + \phi_{Ab} + \phi_{aB} + \phi_{ab})e^{\alpha}(\phi_{AB}e^{\beta} + \phi_{Ab} + \phi_{aB} + \phi_{ab})} \\
&= \beta
\end{aligned}$$

For the *trans* effect

$$logit(p) = \alpha + \beta \cdot Trans$$

the diplotypes have penetrance under the rare disease assumption:

| $f_{ijkl}$ | AB | Ab | aB | ab |
|---|---|---|---|---|
| AB | $e^{\alpha+2\beta}$ | $e^{\alpha+\beta}$ | $e^{\alpha+\beta}$ | $e^{\alpha}$ |
| Ab | $e^{\alpha+\beta}$ | $e^{\alpha}$ | $e^{\alpha+\beta}$ | $e^{\alpha}$ |
| aB | $e^{\alpha+\beta}$ | $e^{\alpha+\beta}$ | $e^{\alpha}$ | $e^{\alpha}$ |
| ab | $e^{\alpha}$ | $e^{\alpha}$ | $e^{\alpha}$ | $e^{\alpha}$ |

Then log odds-ratio

$$\begin{aligned}
log(OR) &= log\frac{h_{AB}h_{ab}}{h_{aB}h_{Ab}} \\
&= log\frac{e^{\alpha}(\phi_{AB}e^{2\beta}+\phi_{Ab}e^{\beta}+\phi_{aB}e^{\beta}+\phi_{ab})e^{\alpha}(\phi_{AB}+\phi_{Ab}+\phi_{aB}+\phi_{ab})}{e^{\alpha}(\phi_{AB}e^{\beta}+\phi_{Ab}+\phi_{aB}e^{\beta}+\phi_{ab})e^{\alpha}(\phi_{AB}e^{\beta}+\phi_{Ab}e^{\beta}+\phi_{aB}+\phi_{ab})} \\
&= log\frac{(\phi_{AB}+\phi_{Ab}+\phi_{aB}+\phi_{ab})(\phi_{AB}e^{2\beta}+\phi_{Ab}e^{\beta}+\phi_{aB}e^{\beta}+\phi_{ab})}{(\phi_{AB}+\phi_{Ab}+\phi_{aB}+\phi_{ab})(\phi_{AB}e^{2\beta}+\phi_{Ab}e^{\beta}+\phi_{aB}e^{\beta}+\phi_{ab})} \\
&= log(1) = 0
\end{aligned}$$

Therefore, the haplotype frequency OR approximates the additive *cis* effects, and cannot capture *trans* effects.
