## Supplementary material for "On the analysis of genetic association with long-read sequencing data": S2 Text

### S2 Text: The coefficients of phase effects for different phasetypes

The choice of reference alleles affects the estimated coefficients for phase effects. Specifically, $\hat{\beta}_{cis}$ for $P_{cis} = Cis_{AB}$ is identical to the one when $P_{cis} = Cis_{ab}$, and opposite in sign to those when $P_{cis} = Cis_{aB}$ and $P_{cis} = Cis_{Ab}$. It is due to the linear relationships between the phasetypes (S1 Text). For example, consider an additive *cis* effect contributed by $Cis_{ab}$:

$$g(E(Y)) = \beta_0^{ab} + \beta_{G_A}^{ab} G_A + \beta_{G_B}^{ab} G_B + \beta_{cis}^{ab} Cis_{ab}$$

$\beta^{ij}$ corresponds to the coefficients with $P_{cis} = Cis_{ij}$. Replace $Cis_{ab}$ with $Cis_{ab} = 2 - G_A - G_B + Cis_{AB}$ (S1 Text), we have:

$$\begin{aligned} g(E(Y)) =& \beta_0^{ab} + \beta_{G_A}^{ab} G_A + \beta_{G_B}^{ab} G_B + \beta_{cis}^{ab} Cis_{ab} \\ =& \beta_0^{ab} + \beta_{G_A}^{ab} G_A + \beta_{G_B}^{ab} G_B + \beta_{cis}^{ab}(2 - G_A - G_B + Cis_{AB}) \\ =& (\beta_0^{ab} + 2\beta_{cis}^{ab}) + (\beta_{G_A}^{ab} - \beta_{cis}^{ab}) G_A + (\beta_{G_B}^{ab} - \beta_{cis}^{ab}) G_B + \beta_{cis}^{ab} Cis_{AB} \\ =& \beta_0^{AB} + \beta_{G_A}^{AB} G_A + \beta_{G_B}^{AB} G_B + \beta_{cis}^{AB} Cis_{AB} \end{aligned}$$

Therefore,

$$\beta_0^{AB} = \beta_0^{ab} + 2\beta_{cis}^{ab};\ \beta_{G_A}^{AB} = \beta_{G_A}^{ab} - \beta_{cis}^{ab};\ \beta_{G_B}^{AB} = \beta_{G_B}^{ab} - \beta_{cis}^{ab};\ \boldsymbol{\beta_{cis}^{ab} = \beta_{cis}^{AB}}$$

We can also show

$$\beta_0^{AB} = \beta_0^{Ab};\ \beta_{G_A}^{AB} = \beta_{G_A}^{Ab} + \beta_{cis}^{Ab};\ \beta_{G_B}^{AB} = \beta_{G_B}^{Ab};\ \boldsymbol{\beta_{cis}^{AB} = -\beta_{cis}^{Ab}}$$

$$\beta_0^{AB} = \beta_0^{aB};\ \beta_{G_A}^{AB} = \beta_{G_A}^{aB};\ \beta_{G_B}^{AB} = \beta_{G_B}^{aB} + \beta_{cis}^{aB};\ \boldsymbol{\beta_{cis}^{AB} = -\beta_{cis}^{aB}}$$

The predicted value $\hat{Y}$ remains unchanged for different reference alleles. For example, $P_{cis} = Cis_{AB}$ and $P_{cis} = Cis_{ab}$ lead to identical $\hat{Y}$, regardless of the presence of marginal effects. It is because the difference between the two phasetypes is fully absorbed through the re-parameterization of the model coefficients. To be specific, consider the design matrices for the linear regression models:

$$X_{AB} = [\mathbf{1}, \boldsymbol{G_A}, \boldsymbol{G_B}, \boldsymbol{Cis_{AB}}] \quad \text{and} \quad X_{ab} = [\mathbf{1}, \boldsymbol{G_A}, \boldsymbol{G_B}, \boldsymbol{Cis_{ab}}]$$

which has the following linear relationships:

$$X_{ab} = X_{AB} \cdot T \quad \text{and} \quad T = \begin{bmatrix} 1 & 0 & 0 & 2 \\ 0 & 1 & 0 & -1 \\ 0 & 0 & 1 & -1 \\ 0 & 0 & 0 & 1 \end{bmatrix}$$

Then the predicted value $\hat{Y}_{ab}$ with $P_{cis} = Cis_{ab}$:

$$\begin{aligned} \hat{Y}_{ab} &= X_{ab}(X'_{ab}X_{ab})^{-1}X'_{ab}Y \\ &= X_{AB}T(T'X'_{AB}X_{AB}T)^{-1}T'X'_{AB}Y \\ &= X_{AB}TT^{-1}(X'_{AB}X_{AB})^{-1}(TT^{-1})'X'_{AB}Y \\ &= X_{AB}(X'_{AB}X_{AB})^{-1}X'_{AB}Y \\ &= \hat{Y}_{AB} \end{aligned}$$

and it is easy to show that $\hat{Y}_{AB} = \hat{Y}_{Ab} = \hat{Y}_{aB} = \hat{Y}_{ab}$.

The same property can be derived for *trans* relationships.
