## Supplementary material for "On the analysis of genetic association with long-read sequencing data": S2 Table

**S2 Table. Impact of phasing uncertainty on RoP.** Switch errors were introduced only in individuals heterozygous at both variants at rates of 5%–20%, with 0% as the benchmark (no phasing error). Haplotypes were simulated for n=5,000 individuals under an additive *cis* effect with heritability $h^2 = 0.003$. Power for detecting *cis* effects and type I error (T1E) for *trans* effects were estimated from 2,000 iterations per error level under three MAF scenarios: Rare-Rare (MAF=0.05 for both alleles), Rare-Common (MAF = 0.05 abd 0.2), and Common–Common (MAF=0.2 for both alleles). RoP retained over 95% of its original *cis* power and maintained nominal *trans* T1E up to 10% switch error, with substantial T1E inflation at 20%.

| **Switch Error Rate** | **Rare–Rare** | | **Rare–Common** | | **Common–Common** | |
|---|---|---|---|---|---|---|
| | **Power (*cis*)** | **T1E (*trans*)** | **Power (*cis*)** | **T1E (*trans*)** | **Power (*cis*)** | **T1E (*trans*)** |
| 0% | 0.973 | 0.052 | 0.950 | 0.044 | 0.925 | 0.049 |
| 5% | 0.973 | 0.046 | 0.943 | 0.051 | 0.891 | 0.056 |
| 10% | 0.936 | 0.060 | 0.916 | 0.058 | 0.876 | 0.059 |
| 20% | 0.886 | 0.122 | 0.841 | 0.106 | 0.802 | 0.110 |
