## Supplementary material for "On the analysis of genetic association with long-read sequencing data": S1 Table

**S1 Table. Summary of simulation results for type I error.** "Yes" corresponds to the correct type I error rate.

| | RoP | Interaction 4 df and Saturated | Interaction 1 df | OR |
|---|---|---|---|---|
| *D'=0* | | | | |
| Model 1: Additive effect on locus A | Yes | Yes | Yes | Yes |
| Model 2: Dominant effect on locus A | Yes | Yes | No | Yes |
| Model 3: Recessive effect on locus A | Yes | Yes | Yes | Yes |
| Model 4: Additive effects on both loci | Yes | Yes | Yes | No |
| Model 5: Dominant effects on both loci | Yes | Yes | Yes | No |
| Model 6: Recessive effects on both loci | Yes | Yes | Yes | No |
| *D'=0.8* | | | | |
| Model 1: Additive effect on locus A | Yes | Yes | Yes | Yes |
| Model 2: Dominant effect on locus A | Yes | Yes | No | Yes |
| Model 3: Recessive effect on locus A | Yes | Yes | No | Yes |
| Model 4: Additive effects on both loci | Yes | Yes | Yes | No |
| Model 5: Dominant effects on both loci | Yes | Yes | No | No |
| Model 6: Recessive effects on both loci | Yes | Yes | No | No |
