## Supplementary material for "On the analysis of genetic association with long-read sequencing data": S1 Text

### S1 Text: Linear relationships between genotypes and phasetypes

Here we show the linear relationships between $Cis_{AB}$ with $Cis_{Ab}$, $Cis_{aB}$ and $Cis_{ab}$, as defined in Table 1 in main text.

$$\begin{aligned} Cis_{ab} &= \begin{bmatrix} 2 & -1 & 0 & -1 & 0 & 0 \end{bmatrix} \cdot \begin{bmatrix} 1 \\ G_A \\ D_A \\ G_B \\ D_B \\ Trans_{AB} \end{bmatrix} + 1 \cdot Cis_{AB} \\ &= 2 - G_A - G_B + Cis_{AB} \end{aligned}$$

$$\begin{aligned} Cis_{Ab} &= \begin{bmatrix} 0 & 1 & 0 & 0 & 0 & 0 \end{bmatrix} \cdot \begin{bmatrix} 1 \\ G_A \\ D_A \\ G_B \\ D_B \\ Trans_{AB} \end{bmatrix} + (-1) \cdot Cis_{AB} \\ &= G_A - Cis_{AB} \end{aligned}$$

$$\begin{aligned} Cis_{aB} &= \begin{bmatrix} 0 & 0 & 0 & 1 & 0 & 0 \end{bmatrix} \cdot \begin{bmatrix} 1 \\ G_A \\ D_A \\ G_B \\ D_B \\ Trans_{AB} \end{bmatrix} + (-1) \cdot Cis_{AB} \\ &= G_B - Cis_{AB} \end{aligned}$$

where $G_A$, $G_B$, $D_A$ and $D_B$ are additive genotypes and the dominance terms (Table 1 main text).

Therefore, testing *cis* effect with $P_{cis} = Cis_{AB}$ is equivalence to $P_{cis} = Cis_{Ab}$, $P_{cis} = Cis_{aB}$ or $P_{cis} = Cis_{ab}$.

The *trans* effects have the same relationships:

$$\begin{aligned} Trans_{ab} &= \begin{bmatrix} 2 & -1 & 0 & -1 & 0 & 0 \end{bmatrix} \cdot \begin{bmatrix} 1 \\ G_A \\ D_A \\ G_B \\ D_B \\ Cis_{AB} \end{bmatrix} + 1 \cdot Trans_{AB} \\ &= 2 - G_A - G_B + Trans_{AB} \end{aligned}$$

$$\begin{aligned} Trans_{Ab} &= \begin{bmatrix} 0 & 1 & 0 & 0 & 0 & 0 \end{bmatrix} \cdot \begin{bmatrix} 1 \\ G_A \\ D_A \\ G_B \\ D_B \\ Cis_{AB} \end{bmatrix} + (-1) \cdot Trans_{AB} \\ &= G_A - Trans_{AB} \end{aligned}$$

$$
\begin{aligned}
Trans_{aB} &= \begin{bmatrix} 0 & 0 & 0 & 1 & 0 & 0 \end{bmatrix} \cdot \begin{bmatrix} 1 \\ G_A \\ D_A \\ G_B \\ D_B \\ Cis_{AB} \end{bmatrix} + (-1) \cdot Trans_{AB} \\
&= G_B - Trans_{AB}
\end{aligned}
$$
